# Hsp10: From Single to Double Rings—Structural Basis of Protein Homeostasis

**DOI:** 10.64898/2026.05.12.724654

**Authors:** Abigail Page, Wyatt Hendricks, Aranea Dunckley, Marielle A. Wälti

## Abstract

The human chaperone complex Hsp60/Hsp10 is essential for maintaining cellular proteostasis by preventing protein misfolding and aggregation. Disruption of these processes contributes to neurodegenerative diseases, while overexpression of Hsp60 and Hsp10 is associated with various cancers. Understanding their molecular mechanisms is therefore of fundamental importance. Unlike its bacterial homolog GroEL, human Hsp60 adopts multiple oligomeric states, with both heptameric and tetradecameric forms binding Hsp10 to form a folding cavity for substrate refolding. Here, we determine the cryo-EM structure of apo Hsp10 and find that, in addition to its single-ring form, it also assembles into a compact double-ring state. This reveals that Hsp10, like Hsp60, exhibits structural behavior that differs markedly from its bacterial counterpart. Traditionally viewed as a passive cofactor assisting Hsp60, we show that Hsp10 alone possesses intrinsic chaperone activity: it supports folding of natural substrates such as malate dehydrogenase 1 and manganese superoxide dismutase. NMR analysis further shows that substrate binding occurs primarily at the core of Hsp10 rather than at the loops. Our findings suggest that Hsp10 exists in equilibrium between single- and double-ring complexes in the unbound state, and upon binding as a single-ring complex, it actively guides substrates into the folding chamber.

---

Molecular chaperones play a pivotal role in assisting nascent polypeptides in achieving their native conformations, preventing aggregation, and facilitating disassembly. The Hsp family comprises several proteins, for example, Hsp60^1^, Hsp70^2^, and Hsp90^3^, named based on their monomeric molecular weights. Although Hsps are expressed under normal physiological conditions, their expression levels are significantly upregulated during cellular stress, such as infection and cancer. This overexpression underscores their cytoprotective and immunoregulatory roles as well as their potential as therapeutic targets^4^. Human Hsp60, encoded by the HSPD1 gene, is essential for protein folding and cellular function, with its inhibition proving detrimental to cell viability^5^. Together with Hsp70, the Hsp60/Hsp10 chaperonin system represents one of the most essential mitochondrial chaperone machineries. Impairment of these processes can lead to severe pathologies, such as protein misfolding and amyloidogenic disorders, for example Alzheimer’s^6,7^, Parkinson’s^8^, and Type 2 Diabetes^9^, while overexpression of Hsp60 and Hsp10 has been observed in various cancers^10-12^. Particular high expression levels are found in cervical, bowel, uterine, and ovarian carcinomas^13-16^ and a recent paper highlights Hsp10 as an actionable cancer treatment target^17^. Elucidating the molecular mechanisms underlying Hsp10 function is of significant biological and medical importance. Studies of the bacterial homolog GroEL/GroES have established a well-defined tetradecameric architecture and folding cycle^18,19^: substrates bind the apical domain of GroEL, ATP binding induces a large conformational change, and GroES association displaces the substrate into the central cavity for folding^1,19-21^. Human Hsp60 is more structurally flexible, existing as monomers, heptamers, or tetradecamers^22,23^, but is thought to follow a similar cycle. In the presence of ATP, both heptameric and tetradecameric forms bind Hsp10 to form a closed chamber that promotes substrate refolding^24-29^. ATP hydrolysis then triggers Hsp10 and substrate release, completing the cycle. However, current structural and functional data remain insufficient to define the conformational transitions from initial substrate binding to encapsulation, leaving unresolved how unfolded proteins are translocated into the chaperonin chamber. Hsp10 has traditionally been regarded as a passive co-chaperonin assisting Hsp60; however, accumulating evidence suggests that Hsp10 may play a more active role in inhibiting amyloid fibril formation^30,31^. The hypothesis that Hsp10 has an independent function is further supported by mRNA data from *The Human Protein Atlas* (www.proteinatlas.org), showing that Hsp10 is present at higher levels than Hsp60, indicating a potential role beyond its interaction with Hsp60.

Here, we show that Hsp10 is in equilibrium between a single and a stable double-ring complex in solution at physiological concentrations. Using cryo-EM, we determined the structure of the double ring species at ~3.7 Å resolution, and the one of the single ring species at 4 Å resolution. We show that Hsp10 has a function beyond amyloid fibril inhibition. Specifically, we demonstrate that Hsp10 is involved in the folding of native substrates of the Hsp60/10 chaperonin system, including malate dehydrogenase 1 (MDH1) and manganese superoxide dismutase (MnSOD), both key regulators of cellular metabolism. Furthermore, we show that Hsp10 interacts with its natural substrates primarily through its core rather than the loop regions. This observation supports a mechanism in which Hsp10 actively pushes substrates into the Hsp60 folding chamber and thus contributes to protein encapsulation. Our findings indicate that Hsp10 functions not merely as a passive cofactor but as an active participant in maintaining protein homeostasis, directly contributing to substrate encapsulation within the Hsp60 cavity.

## Hsp10 promotes the folding of MDH1 and MnSOD

Hsp10 has previously been shown to inhibit the fibrillization of amyloid-beta 1–42 (Aβ42)^31^ and α-synuclein^30^, intrinsically disordered proteins that aggregates into neurotoxic, β-sheet-rich fibrils found in plaques within the brains of patients with Alzheimer’s disease or Lewy Bodies in Parkinson’s disease. To investigate whether Hsp10 can also assist in substrate folding, we performed refolding assays using the native substrates MDH1 and MnSOD. All proteins for these studies were purified from bacterial overexpression (Supplementary Information). Proper folding of MDH1 was assessed by its ability to catalyze the conversion of NADH to NAD^+^, with the rate of NADH consumption serving as an indicator of successful protein folding. Hsp10 was significantly involved in the folding of MDH1. In the absence of Hsp10, after 170 minutes only approximately 18.5% of the total NADH were converted (Fig. 1a). Whereas in presence of Hsp10 close to 100% NADH was consumed after approximately 160 minutes, similar levels to that observed for MDH1 that was never unfolded (Fig. 1a). Finally, the ability of Hsp10 to refold MnSOD was assessed using two approaches, the first being a commercially available colorimetric superoxide dismutase activity assay. In this assay, xanthine oxidase generates superoxide radicals that react with a colorless substrate to produce a yellow product. Functional MnSOD reduces superoxide accumulation, resulting in decreased absorbance at 450 nm (the yellow product). HCl-denatured MnSOD alone failed to reduce superoxide, as indicated by increased absorbance at 450 nm depicting radical accumulation over time (Fig. 1b). In contrast, in the presence of Hsp10, denatured MnSOD recovered activity comparable to that of folded MnSOD that was not exposed to HCl. Second, far-UV circular dichroism (CD) spectroscopy, which reports on secondary structure content in solution, was used. Native MnSOD contains ~52% α-helical content^32^, whereas Hsp10 consists exclusively of β-strands and lacks α-helices; thus, the α-helical signal detected in the spectra can be attributed solely to MnSOD (Supplementary Fig. 1). MnSOD was first denatured, and refolding was monitored by recording CD spectra in the presence or absence of Hsp10 over 15 minutes. A recovery of ellipticity at 222 nm, indicative of α-helical structure, was observed for denatured MnSOD only in the presence of Hsp10, approaching that of native MnSOD, but not in its absence (Fig. 1c). Before denaturation, MnSOD exhibited 49.6% α-helical content, which decreased to 25.6% upon denaturation. After 15 minutes, denatured MnSOD alone retained 26.6% α-helical content, whereas denatured MnSOD incubated with Hsp10 reached 56% α-helical content. These results indicate that Hsp10 promotes refolding of MnSOD to its native-like state (Fig. 1c and Supplementary Fig. 1). Taken together, these assays demonstrate that Hsp10 promotes the refolding of both MDH1 and MnSOD in the absence of Hsp60, indicating that it is not merely a passive co-chaperone but can act independently to guide substrate folding. This intrinsic refolding activity suggests that Hsp10 possesses chaperone-like properties beyond its canonical role as a cap for the Hsp60 folding chamber and supports a model in which Hsp10 actively contributes to maintaining protein homeostasis.

**Fig. 1:**
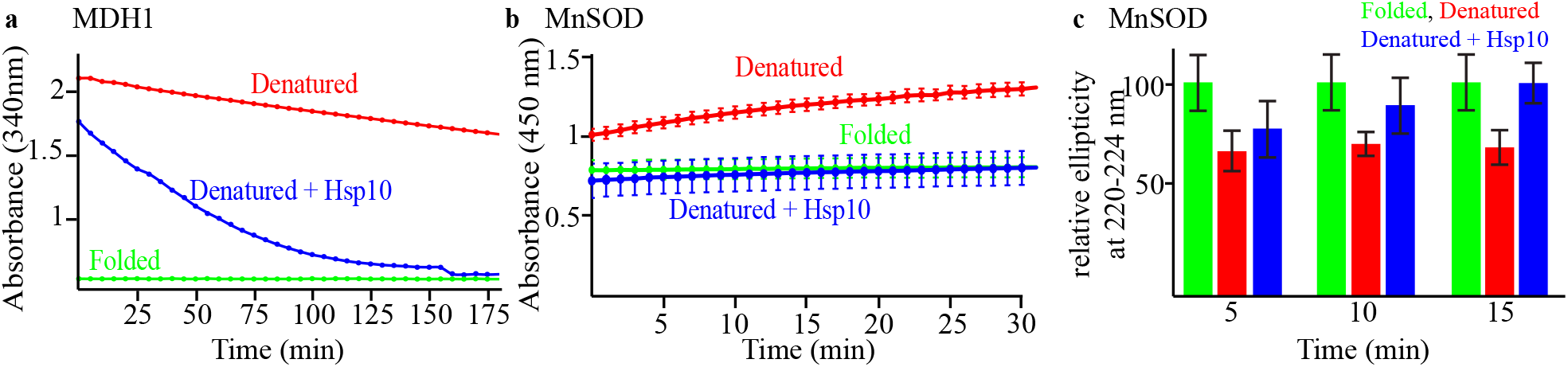
Assessment of Hsp10’s functionality. **a**, MDH1 time-course activity assay. The folding of MDH1 was assessed by NADH depletion in absence (red) and presence of Hsp10 (blue), folded MDH1 is shown in green. **b**, Time course of SOD activity. Superoxide radical production was measured at 450 nm over time with denatured MnSOD in the absence (red) and presence (blue) of Hsp10. Never denatured MnSOD is shown in green. **c**, average ellipticity at 220-224 nm Folding of MnSOD relative to the folded sample (green) was assessed using far-UV circular dichroism over a 15-minute timespan. Denatured MnSOD in absence (red) and presence (blue) of Hsp10. Error bars indicate the standard deviation.

### In solution, Hsp10 exists in equilibrium between single- and double-ring species

The bacterial co-chaperone GroES forms a single heptameric ring (Fig. 2a,b)^33^. In contrast, Hsp10 assembles into both heptameric single-ring and tetradecameric double-ring complexes at concentrations above 10 nM, as observed by native gel electrophoresis (Fig. 2a) and mass photometry (Fig. 2c-j). For mass photometry, accurate detection requires that no other particles land within the spatial area defined by the diffraction-limited spot or within the temporal window set by the integration time. Any additional landing events in close proximity—either in space or time—can distort the signal and compromise data quality. Consequently, to minimize coincident events, measurements are typically restricted to low analyte concentrations, generally on the order of a few tens of nanomolar. For higher concentration measurements the cover slides were treated by amination and PEGylation to lower the surface binding, according to a previously published protocol^34^. At 10 μM Hsp10, approximately half of the molecules populate a single-ring species (42%) and 53% a double-ring species. However, increasing the salt concentration to 1 M NaCl or lowering the pH to 5 shifts the equilibrium to favor the single-ring species, with its population rising to over 70% at high salt and to about two-thirds at low pH (Fig. 2c, and 2d). The populations are concentration dependent: at 1 nM, a fully heptameric species was observed, whereas at 100 μM, Hsp10 is predominantly tetradecameric (Fig. 2f–k). Using Blue Native Polyacrylamide Gel Electrophoresis (BN-PAGE), we show that Hsp10 retains its canonical function as a cap that closes the barrel-shaped Hsp60 complex. Hsp10 was incubated with Hsp60 in the presence of ATP for 10 minutes at 30 °C. Mizoribine^35^, a known inhibitor of Hsp60 ATPase activity, was added prior to loading the samples onto the BN gel. Distinct bands were observed corresponding to Hsp10 binding to both the tetradecameric and heptameric forms of Hsp60, resulting in the formation of the so-called “football” (Hsp10_7_–Hsp60_14_–Hsp10_7_) and “half-football” (Hsp10_7_–Hsp60_7_) complexes, respectively (Supplementary Fig. 2). We do not observe any evidence of Hsp10 remaining in a double-ring species in the presence of Hsp60 and ATP.

**Fig. 2:**
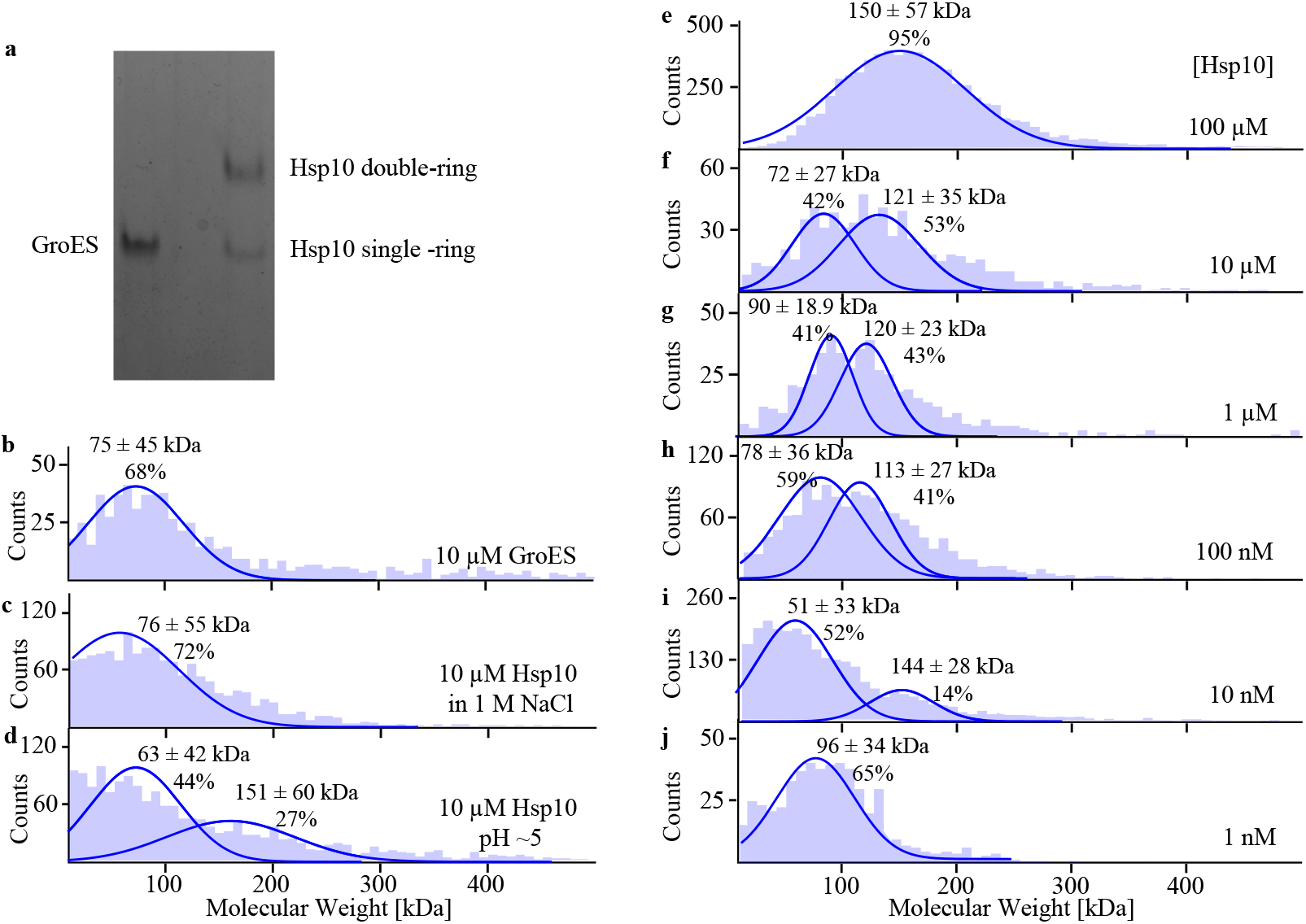
Size distribution of GroES and Hsp10. **a**, BN–PAGE of 10 μM GroES and 10 μM Hsp10. **b**, Mass photometry analysis of GroES. **c**, Oligomeric state of 10 μM Hsp10 in presence of 1 M NaCl or **d**, at pH 5. **e–j**, Oligomeric state of Hsp10 measured at the indicated concentrations. Samples were recorded at room temperature in 20 mM KCl, 20 mM magnesium acetate, 50 mM Tris–HCl (pH 7.4), 300 mM NaCl, and 10 mM MgCl_2_, unless otherwise stated.

### Cryo-EM structure of Hsp10

Two samples were prepared for cryo-electron microscopy (cryo-EM): one containing Hsp10 at 8 mg/ml and another at 2 mg/ml. In the higher-concentration sample, the equilibrium was strongly shifted toward the double-ring structure, with approximately 32-fold more particles in the double-ring conformation than in the single-ring state. In contrast, the lower-concentration sample displayed an approximate 1:1 ratio of double- and single-ring particles.

A total of 137,564 particles were used to determine the structure of the double-ring Hsp10 complex at an overall resolution of ~3.7 Å, applying C7 symmetry, with local resolution ranging from 2.5 to 4.0 Å. (Fig. 3, Supplementary Fig. 3, and Supplementary Table 1). Approximately 33% of the particles adopt a top view, 57% a side view, and 10% a corner view. The double ring adopts a back-to-back arrangement of two single rings. The structure comprises six well-resolved β-strands in each ring (Supplementary Fig. 4), stabilized by multiple hydrogen bonds. Within each ring, β-strands from adjacent subunits form a continuous sheet composed of β2–β3–β4–S2–β1–S1. These interactions are also observed in the bound Hsp10 structure (PDB 6MRC) (Supplementary Fig. 5). In addition to these strands, two short β-strand–like segments comprising residues 5–6 and 98–99 are observed and are stabilized by hydrogen bonding. The two rings adopt distinct conformations, indicating structural asymmetry within the complex. In particular, β-strand 2 (residues 15–20) and β-strand 6 (residues 87–91) are laterally offset between the two rings (Supplementary Fig. 6). The rings are held together by hydrophobic interactions involving residues A5, V16, L17, L90, and F91. In addition, salt bridges involving Arg15 and Arg93 stabilize the double-ring assembly, while Arg92 forms stabilizing interactions with Asp93 (Fig. 3c). Residues 1–4, 7–8, 12–14, 21–42, 50–70, 74–78, 84–86, 92–97, and 100–102 were not resolved in the electron density map, suggesting that these regions are highly flexible. These segments include the N- and C-terminal residues, disordered regions, and two large mobile loops (residues 21–42 and 50–70).

**Fig. 3:**
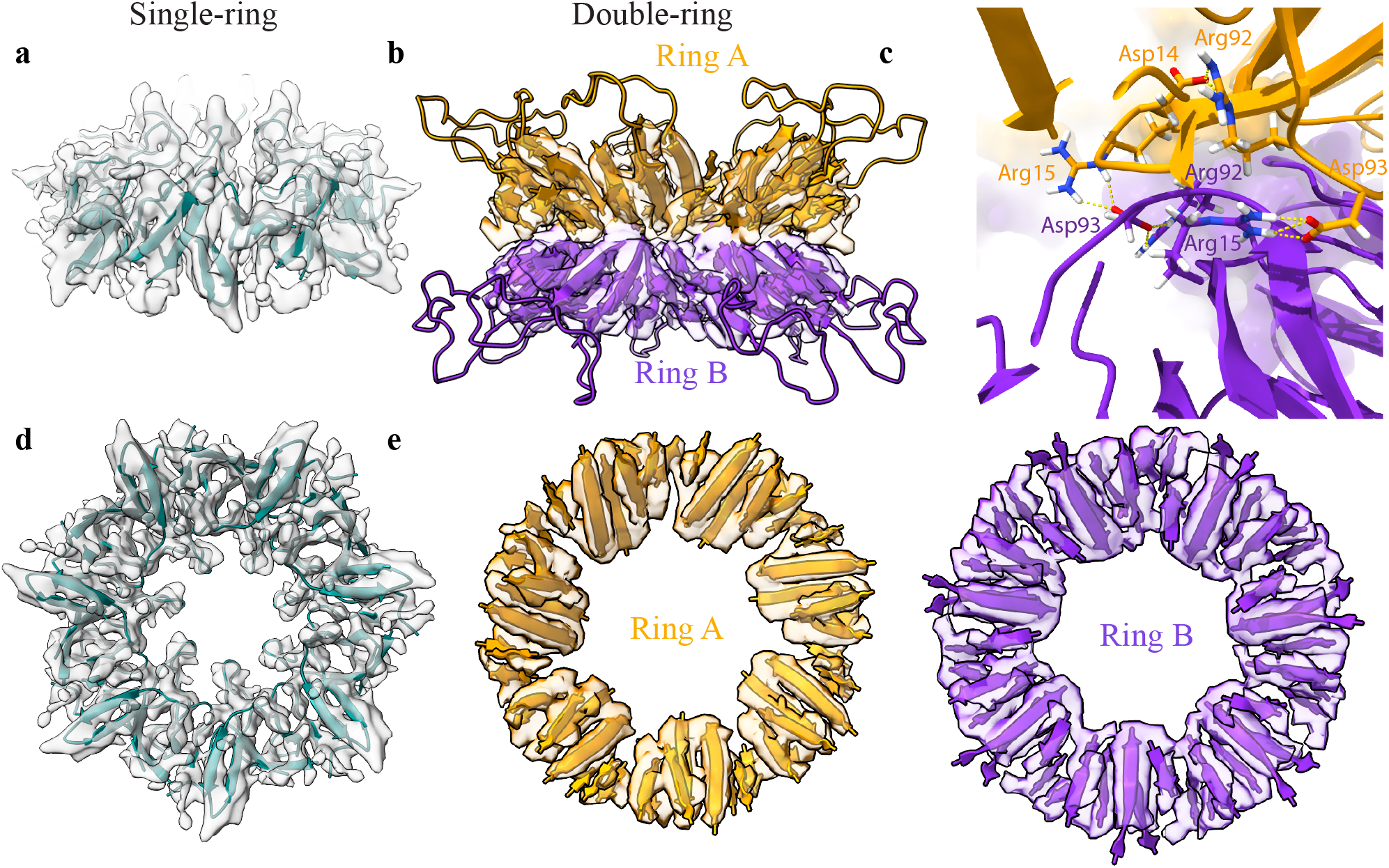
Cryo-EM structure of the single- and double-ring Hsp10 complex. Electron density maps with corresponding atomic models are shown as side views for **a**, single-ring Hsp10 and **b**, double-ring Hsp10, in which rings A (orange) and B (purple) are arranged back-to-back. Loops corresponding to residues 20–42 are modeled. **c**, Inter-ring ionic bridges are indicated by orange dashed lines. **d**, Bottom views of the electron density maps for the single-ring, and **e**, each double-ring Hsp10 is shown separately with its atomic model.

For the unbound single-ring Hsp10 complex, a total of 46,178 particles were picked from micrographs of grids prepared with a sample concentration of 2 mg/mL (Supplementary Table 1). Due to preferred orientation observed in the micrographs, manual reduction of top-view particles was performed to improve angular sampling and reconstruction quality. In the final particle stack, approximately 24% of particles corresponded to top views, while 76% represented side or tilted orientations. Only residues 1, 2, and 21–42 were not resolved in the electron density map, suggesting that these regions are highly flexible. The final structure was reconstructed using C7 symmetry to an overall resolution of 4 Å (Fig. 3 and Supplementary Fig. 7). Similar to the Hsp10-bound structure and the double-ring complex, the apo single-ring Hsp10 comprises six β-strands (Supplementary Fig. 8). However, distinct structural differences are observed between the bound and unbound conformations (Supplementary Figs. 5 and 9). In the bound state, the β-strands forming the barrel (β2, β3, β5, and β6) are arranged at relatively uniform distances, resulting in a compact and symmetric β-barrel. In contrast, the apo structure shows increased separation between β2–β3 and β5–β6, leading to a more open and less compact barrel architecture (Supplementary Fig. 9). Notably, GroES forms an even more tightly packed β-barrel than the Hsp10-bound conformation^36^.

2D class averages show that top-view projections of the single-ring Hsp10 assembly display more well-defined, higher-resolution features than those of the double-ring species. This is consistent with the greater symmetry and structural homogeneity of the single-ring complex, which facilitates more accurate particle alignment and averaging. In contrast, the double-ring assembly exhibits reduced apparent resolution in top views, likely due to its stacked architecture, increased heterogeneity, and superposition of densities from two asymmetric rings. The central pore is also smaller in the double-ring assembly (~38 Å) compared to the single-ring structure (~45 Å), with additional density extending toward the center from ring B (Supplementary Fig. 10). Side-view class averages further distinguish the two states. The single-ring structure shows a single layer of density with weak peripheral signal corresponding to flexible apical loops (residues 50–70). In contrast, the double-ring assembly displays two parallel density layers consistent with stacked rings, with evidence of inter-ring contacts between adjacent subunits.

When comparing hydrophobic and charged residues within the cavity of Hsp10, we find that the free double-ring conformation is more hydrophobic than the conformation bound to Hsp60 (Fig. 4). Whereas the internal cavity of the double-ring Hsp10 contains 65% (ring A) and 68% (ring B) hydrophobic amino acids, Hsp10 in the bound conformation contains only 47%. In contrast, the cavity of bound Hsp10 contains 35% charged amino acids, whereas the free double-ring only 10% (ring A) and 13% (ring B). The free single-ring Hsp10 also displays increased hydrophobicity in the cavity, with 57% being hydrophobic and 23% being charged. Similarly, in Hsp60 it has been shown that, in the condensed conformation, the inner surface of the cavity is more hydrophobic than in the ATP-bound open conformation, where the surface becomes more negatively charged^1^ and, in humans, more positively charged^27^. This change in the surface properties of the Hsp60 cavity has been suggested to facilitate protein folding by promoting hydrophobic collapse. Our results suggest that Hsp10 may function in a similar manner, with the lid of the cavity exhibiting comparable surface characteristics that could further promote substrate folding.

**Fig. 4:**
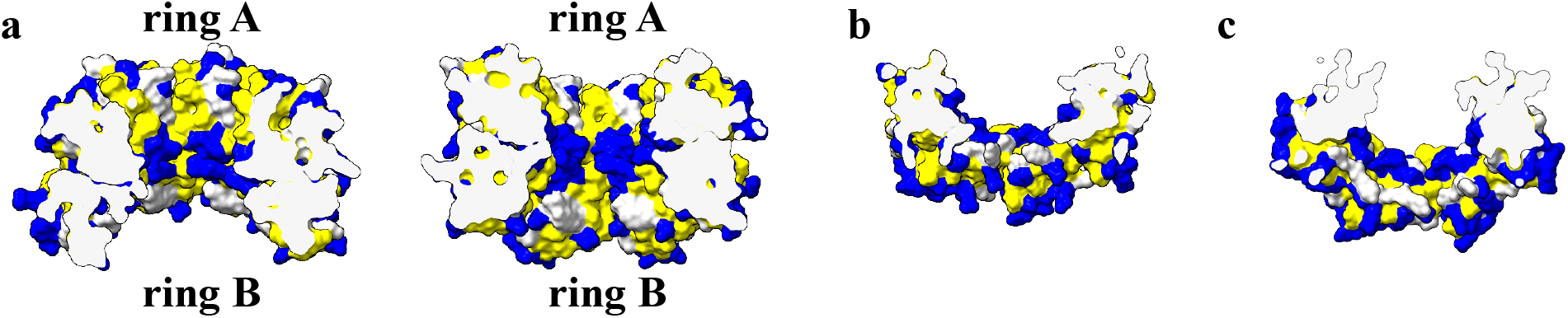
Hydrophobic and charged amino acids highlighted in the core of Hsp10. Hydrophobic residues (yellow) and charged residues (blue) **a**, in the double-ring Hsp10 structure and **b**, in the single-ring free form and **c**, in single ring conformation when bound to Hsp60 (PDB ID: 6MRC)^24^. The loops, residues 21-42, and 53-61, were removed in b and c for comparison.

### Solution NMR only depicts the heptameric species

Solution-state NMR of a deuterated Hsp10 sample shows at least 108 peaks, of which 98 were assigned to the heptameric symmetric Hsp10^37^. The assignment yielded continuous sequence coverage, with signals observed for nearly all residues (95% assigned). Several unassigned peaks were also detected: approximately 10 additional resonances appear as strong peaks, while a larger number of weaker resonances may arise from the flexible loops of the tetradecamer. Based on this high level of assignment completeness, we conclude that the observed NMR signals predominantly arise from the heptameric species. In contrast, the larger asymmetric tetradecameric complex (~140 kDa) does not significantly contribute to the observed spectra due to its large molecular size and the associated line broadening. Comparison of the signal intensity of 90 μM GroES with the same concentration of Hsp10 clearly shows that the intensity of the Hsp10 peaks is around 30% lower than that of GroES (Supplementary Fig. 11). This indicates the loss of a fraction of the sample to the NMR-invisible double-ring species.

### Visualizing the interaction of Hsp10 with natural substrates

Hsp10’s ability to inhibit Aβ42 aggregation has been reported previously^31^, but the structural basis and interaction sites remained unknown. To map the substrate-binding sites on Hsp10, we recorded ^1^H–^15^N HSQC-TROSY spectra of 120 µM ^2^H, ^15^N-labeled Hsp10 in the presence and absence of 400 µM Aβ42, 1 mM MDH1, or 120 µM MnSOD (Supplementary Fig. 12-14). The resulting chemical shift perturbations are shown in Fig. 5. The NMR assignment of Hsp10 was previously deposited in the BMRB (accession no. 53226)^37^. Colloquially, Hsp10 acts as a cap to close the cavity formed by Hsp60, capturing substrates within this chamber for folding. The “inner side” of Hsp10 refers to the surface facing the internal cavity of the Hsp60/Hsp10 complex after closure. In the presence of Aβ42, chemical shift perturbations were observed across multiple residues spanning nearly the entire Hsp10 structure, indicating widespread interaction (Fig. 5a). In contrast, binding of MDH1 and MnSOD to Hsp10 produced more localized perturbations, confined to defined regions (Fig. 5b,c). While a small subset of interactions occurred at the flexible loop regions of Hsp10, 6 residue for MDH1 (out of 31 total) and 3 residues for MnSOD (out of 22 total), the majority of interactions were localized to the structured core surrounding the β-barrel–like region of Hsp10. Together, these data indicate that Hsp10 serves as an additional binding platform for substrate proteins alongside Hsp60. Notably, different client proteins preferentially interact with the β-core rather than the flexible loop regions of Hsp10 supporting a model in which Hsp10 plays an active role in substrate encapsulation.

**Fig. 5.**
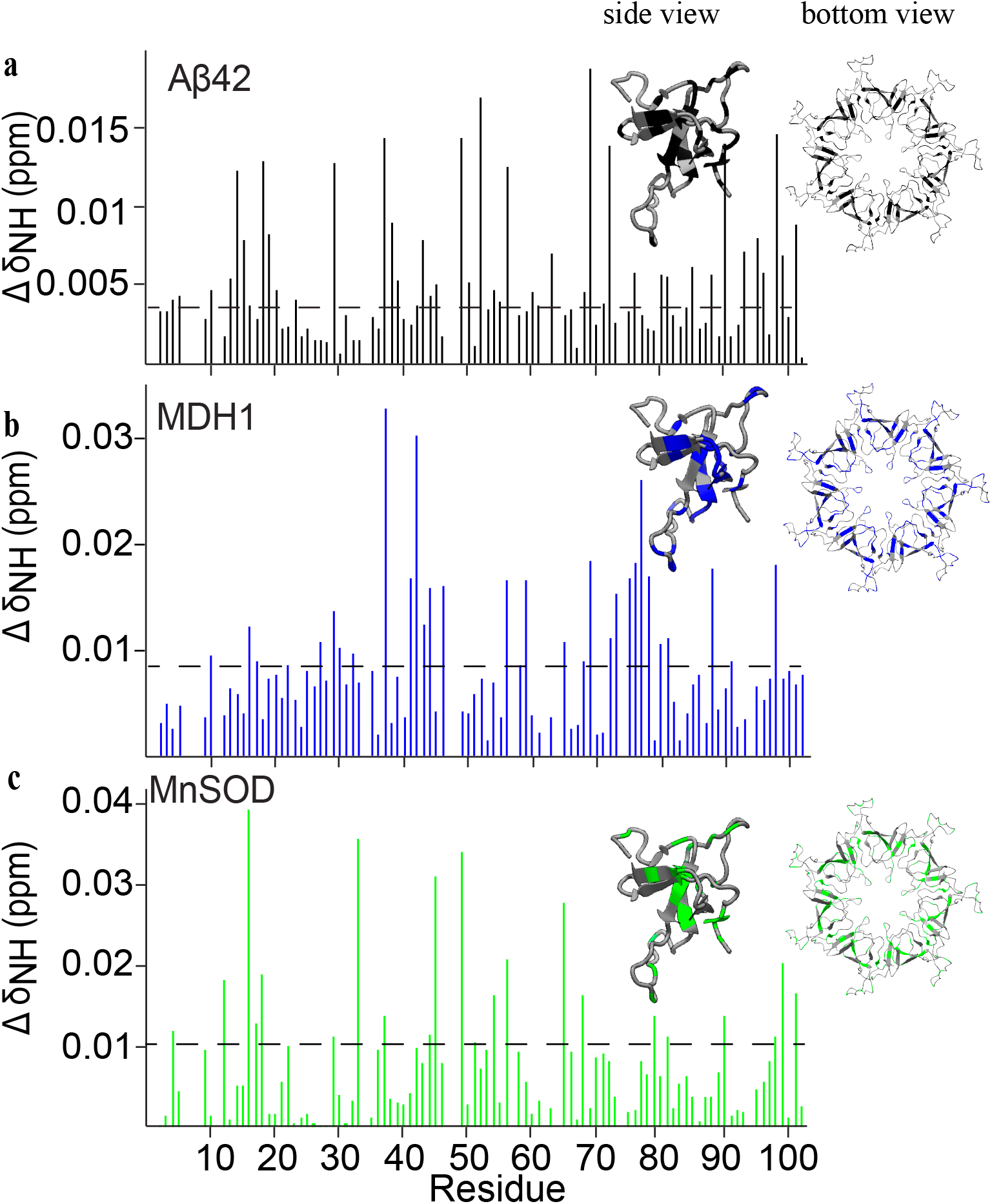
Interaction of substrates with the single-ring Hsp10. Chemical shift perturbation (ΔδNH) of 120 μM [^2^H,^15^N]-labelled Hsp10 upon the addition of **a**, 500 μM Aβ42 at 277 K, **b**, 1 mM MDH1 at 298 K, and **c**, 120 μM MnSOD at 298 K. Chemical shift perturbation was calculated using the formula [(0.14ΔδN^2^+ΔδH^2^)/2]^1/2^, as described previously^38^. The threshold was defined according to Williamson^38^ and was set to 3σ for MnSOD, 4σ for MDH1, and 5σ for Aβ42. The corresponding interactions are depicted on the single-ring, unbound Hsp10 structure in all insets, with the side view of one subunit shown on the left and the full ring on the right.

**Fig. 6.**
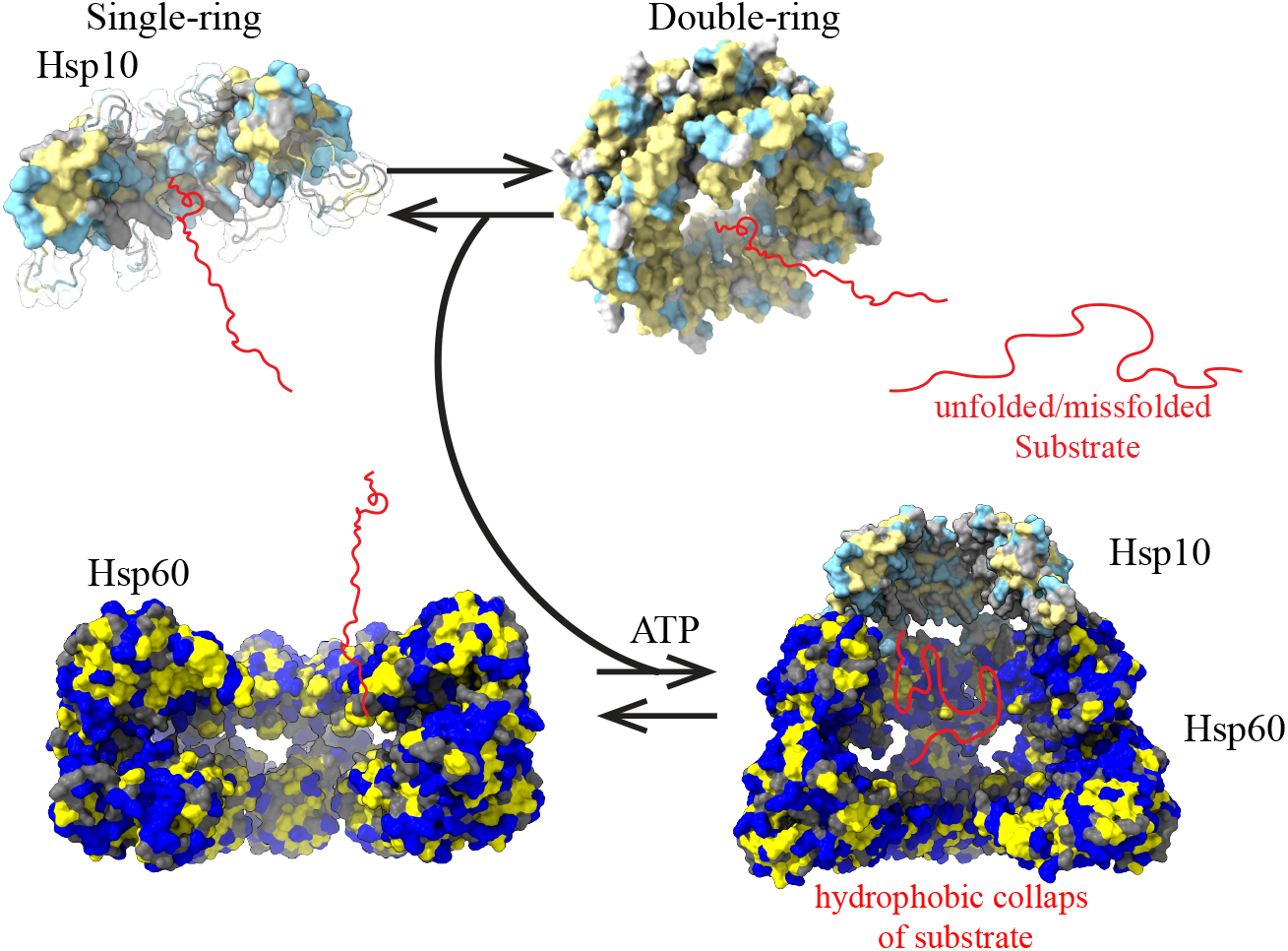
Proposed mechanism of Hsp10 in supporting Hsp60. In solution, Hsp10 exists in an equilibrium between single- and double-ring species. Substrate binding to the apical domain of Hsp60 is further supported by interaction with Hsp10. The double-ring species of Hsp10 exhibits a more hydrophobic cavity surface than the single-ring form and may therefore enhance substrate binding. Upon association with Hsp60, Hsp10 facilitates encapsulation of substrates within the Hsp60 folding chamber.

In this study, we challenge the traditional view of Hsp10 as merely a passive cochaperone and instead highlight its integral role in the coordinated activity of the Hsp10/Hsp60 chaperone machinery. Our data demonstrate that human Hsp10 exists in solution as a dynamic equilibrium between heptameric and tetradecameric assemblies. We solved the structures of both and see that the equilibrium is highly concentration dependent, having higher double-ring Hsp10 species at high concentration. Most residues involved in oligomer stabilization, including hydrophobic contacts and salt bridges between Arg15 and Asp93, are further stabilized by an internal salt bridge with Arg92. GroES contains fewer charged residues in the C-terminal region, and this deficiency may contribute to its inability to form a double-ring species (Supplementary Fig. 15). Beyond its structural properties, our results show that Hsp10 retains functional activity independent of Hsp60. When examining the interaction surfaces, we find that natural substrates such as MDH1 and MnSOD, which likely benefit from encapsulation, primarily bind to the β-core of Hsp10 rather than the loops, leaving the loops available for Hsp60 binding. This observation highlights the importance of the Hsp10 cavity as a conserved substrate-binding site capable of accommodating structurally diverse client proteins. We propose that Hsp10 actively assists in guiding substrates into the folding chamber of Hsp60. The double-ring species may become particularly important under stress conditions, as increased Hsp10 expression is expected to shift the equilibrium toward this state, which exhibits a more hydrophobic inner surface. This property may enhance interactions with unfolded or misfolded substrates. Because substrate binding primarily involves the β-sheet core, the apical loops remain available for interaction with Hsp60. Upon binding to Hsp60, the environment becomes more charged, consistent with a transition to a folding-competent state that facilitates substrate encapsulation and folding. Together, these findings expand the functional understanding of Hsp10, suggesting a more direct role in substrate recognition and delivery within the Hsp10/Hsp60 chaperone system than previously appreciated.

## Additional information

Supplementary Information is available for this paper.

## Acknowledgement

This research was supported by the National Institute of General Medical Sciences of the National Institutes of Health under award number R00GM132496 and R35GM156786. A portion of this research was supported by NIH grant R24GM154185 and performed at the Pacific Northwest Center for Cryo-EM (PNCC) with assistance from Vamseedhar Rayaprolu. We thank Dr. Muhammed Shafeek O.H. (University of Arizona - CBC Molecular Structure Core (MSC) NMR Facility, RRID: SCR_022888) for maintaining the NMR instruments. We thank Professor Thomas Tomasiak for insightful discussions regarding the cryo-EM structure. This work was further supported by the University of Arizona Research, Innovation & Impact (RII) and Technology Research Initiative Fund/Improving Health and Access and Workforce Development.

## Author contributions

A.P. and W.H. prepared the clones, purified the protein, prepared the samples, and performed all measurements. A.D. optimized the MDH1 functional assay. A.P. and M.A.W. recorded the NMR data, and A.P. collected the cryo-EM data at the Pacific Northwest Center for Cryo-EM

(PNCC) with their support. A.P., W.H., and M.A.W. analyzed all the data. A.P. and M.A.W. wrote the manuscript. All authors reviewed the manuscript.

## Data availability

The atomic coordinates and electron micrographs for the Hsp10 single-ring species have been deposited in the Protein Data Bank (PDB) and Electron Microscopy Data Bank (EMDB) under accession codes 13KW and EMD-77126, respectively. The Hsp10 double-ring species has been deposited under accession codes 13KK and EMD-77112, respectively.

