## Supporting Information for "Hsp10: From Single to Double Rings—Structural Basis of Protein Homeostasis"

### Methods

#### TEV protease expression and purification

Purification of the TEV protease was performed as described previously[1]. To summarize, the $His_6$-TEV plasmid was purchased from Addgene (pDZ2087) and transformed into BL21 (DE3) Codon+ RIL cells. This was left to grow overnight in 15 mL Luria-Bertani (LB) at 37°C. The next morning, the preculture was transferred to 1 L of LB supplemented with ampicillin and chloramphenicol and let to grow to an $OD_{600}$ of 1.2. After induction with 0.5 mM isopropyl-B-D-thiogalactoside (IPTG), the temperature was lowered to 30°C and the cells were left to express overnight. The next morning, the cells were centrifuged at 3,500 xg for 25 minutes at 4 °C. The subsequent pellet was resuspended with 50 mL of 50 mM Tris, pH 8.0, 200 mM NaCl, 20 mM imidazole, and 10% v/v glycerol, 0.4 g of streptomycin sulfate and 1 tablet of cOmplete Protease Inhibitor Cocktail tablet. After 30 minutes, the sample was ran 4x through an Avestin emulsifier to emulsify the cells. The lysate was then centrifuged at 23,500 xg for 25 minutes at 4°C. The supernatant was loaded onto a 5 mL nickel column (HisTrap™, HP, GE Healthcare) and eluted with varying levels of 50 mL of 50 mM Tris, pH 8.0, 200 mM NaCl, 1 M imidazole, and 10% v/v glycerol. The fraction containing the TEV protease was concentrated and loaded onto a Superdex™ 75 (Cytiva) size exclusion column equilibrated with 50 mM Tris, pH 8.0, 200 mM NaCl, 10% v/v glycerol, and 2 mM EDTA. SDS-PAGE gel electrophoresis was used to confirm purity. The TEV protease containing fractions were concentrated to 1 mg/mL and stored at -20 °C.

#### Expression and Purification of Hsp10

The human cpn10 gene, cloned into the pET21d plasmid, was ordered from GenScript with codon optimization and an N-terminal 6×His tag, a GST tag, and a TEV protease cleavage site. The plasmid was transformed into *E. coli* BL21(DE3) cells amplified in house.

For unlabeled Hsp10, 1 colony from a transformation was growth in 15 mL of LB supplementef with ampicillin and choroamphenicol. This growth was transferred into 1 L the next morning and the cells were grown at 37 °C in LB to an $OD_{600}$ of 0.8–1.2, induced with 0.5 mM IPTG, shifted to 20 °C, and left to expressed overnight.

For $^{15}N$ $^{12}C$ $^{2}H$-labeled Hsp10 and $^{15}N$ $^{13}C$ $^{2}H$-labeled Hsp10, all colonies from one agar plate were used to inoculate 200 mL LB and grown at 37 °C overnight. The following morning, cells were harvested by centrifugation at 3,500 xg for 10 minutes, and the pellet was resuspended in 1 L deuterated minimal medium ($^{2}H$-M9) containing either $^{15}N$ ammonium chloride and unlabeled glucose or $^{15}N$ ammonium chloride and $^{13}C$ d7 glucose.[2] Protein expression was induced and carried out under the same conditions as for unlabeled Hsp10.

Cells were harvested by centrifugation at 3,500 xg for 25 minutes at 4 °C. The pellet was lysed using an Avestin Emulsiflex in 40 mL lysis buffer (10 mM imidazole, 300 mM NaCl, 50 mM Tris-HCl, pH 8.0) supplemented with 0.4 g streptomycin sulfate and one cOmplete™ EDTA-free Protease Inhibitor Cocktail tablet (Roche, MilliporeSigma). The lysate was clarified by centrifugation at 23,500 xg for 25 minutes at 4 °C, after which the supernatant was incubated again with 0.4 g streptomycin sulfate for 30 minutes at room temperature. The sample was centrifuged at 3,500 xg for an additional 10 minutes, filtered through a 0.45 μm syringe filter, and loaded onto a 5 mL nickel column (HisTrap™, HP, GE Healthcare) pre-equilibrated with 10 mM imidazole, 300 mM NaCl, and 50 mM Tris-HCl at pH 8.0 at a flow rate of 2 mL/min. After washing with 6% and 25% buffer B (1 M imidazole, 300 mM NaCl, 50 mM Tris-HCl, pH 8.0) to remove non-specific binding to the column, Hsp10 was eluted with 100% buffer B. The eluted protein was dialyzed against 1 mM EDTA, 10 mM imidazole, 50 mM Tris-HCl at pH 7.4, 200 mM NaCl, and incubated overnight at room temperature for tag cleavage. TEV protease was added prior to dialysis at a 1:25 ratio of Hsp10 to TEV. Following dialysis, $MgCl_2$ was added to quench the EDTA prior to loading to the column, and the sample was loaded onto a second 5 mL nickel column. The flow-through was concentrated using 30 kDa molecular weight cutoff Vivaspin® Turbo Regenerated Cellulose Centrifugal Concentrators (Sartorius). Protein purity was assessed by SDS-PAGE, and the oligomeric state was analyzed by Blue Native PAGE (Invitrogen). If not immediately used the sample is concentrated and frozen at -20 °C at ~500 μM.

**Hsp10 with a Thrombin cleavage site**

Hsp10 with a Thrombin cleavage site was expressed and grown as previously stated for the Hsp10 with a TEV cleavage site purification, except with a plasmid containing the CPN10 gene, 6xHis tag, and a Thrombin cleavage site. The lysate was loaded onto a 5 mL nickel column (HisTrap™, HP, GE Healthcare) and eluted from the column with the buffers stated above. After elution, the fraction containing Hsp10 was concentrated and loaded to a 340 mL superdex s75 column (Cytivia) equilibrated with 40 mM sodium phosphate pH 7.0. The final sample was concentrated to 175 µM and frozen at -20ºC. The thrombin cleavage site and 6x his-tag was left on the purified protein to run microscale thermocalorimetry (described later), which required a his-tag on the measured protein.

**Expression and Purification of MtHsp60**

The human mt-cpn60 gene, lacking the disordered 26–amino acid N-terminal mitochondrial import signal, was synthesized by GenScript with codon optimization and cloned for expression with an N-terminal 6×His tag followed by a TEV protease cleavage site. Purification was performed as previously described.[3] The plasmid was transformed into *E. coli* Rosetta cells (amplified in house) which were grown and induced as described previously. Following cell lysis, the lysate was loaded onto a 5 mL nickel affinity column and eluted with 250 mM imidazole. TEV protease was added at a ratio of 1:25 in the protein-containing fraction. The fraction was then dialyzed against 200 mM NaCl 50 mM Tris-HCl pH 7.4 for 1 hour. Following cleavage, the sample was passed over a second 5 mL nickel affinity column, and the flow-through was collected and concentrated to 18–20 mg/mL. A high-concentration buffer was then added to achieve final conditions of 20 mM KCl, 20 mM magnesium acetate, and 10 mM $MgCl_2$. Assembly was induced by the addition of 10% (v/v) glycerol and 10 mM ATP. The sample was incubated overnight at room temperature and subsequently loaded onto a 24 mL Superdex S200 size-exclusion column (citivia) equilibrated with 300 mM NaCl, 50 mM Tris-HCl pH 7.4, 20 mM KCl, 20 mM Magnesium Acetate, and 10 mM $MgCl_2$ to isolate the heptameric and tetrameric fractions. The subsequent fractions were concentrated and left at room temperature for up to a week for further use.

**Purification of MDH1**

Codon-optimized MDH1 cloned into the pET21d plasmid was synthesized by GenScript and contained an N-terminal 6×His tag, a GST tag, and a TEV protease cleavage site. The plasmid was transformed into *E. coli* BL21 cells and grown at 37 °C in LB medium to an $OD_{600}$ of 0.8-1.2. Expression was induced with 0.5 mM IPTG, and cultures were grown overnight at 20 °C. Cells were harvested by centrifugation at 3,500 xg for 20 min, and the cell pellet was resuspended in 40 mL of lysis buffer containing 25 mM Tris-HCl, pH 7.2, 150 mM NaCl, 10 mM imidazole, 5% glycerol, 0.4 g streptomycin sulfate, and one protease inhibitor tablet. Cells were lysed using an Avestin Emulsiflex. The lysate was clarified by centrifugation at 23,500 xg for 25 min at 4 °C. The supernatant was incubated again with 0.4 g streptomycin sulfate and filtered through a 0.45 µm syringe filter before loading onto a 5 mL nickel affinity column. The column was washed sequentially with buffer containing 25 mM Tris-HCl at pH 7.2, 150 mM NaCl, 60 mM imidazole, and 5% glycerol, followed by the same buffer containing 250 mM imidazole. Protein was then eluted with the same buffer containing 1 M imidazole. TEV protease was added to the MDH1-containing fractions at a 1:25 (w/w) ratio and dialyzed against 25 mM Tris-HCl at pH 7.2, 150 mM NaCl, 1 mM β-mercaptoethanol (BME), and 5% glycerol. The sample was then passed over a second nickel affinity column to remove TEV protease, the His-tag, and any uncleaved MDH1 or contaminants. Sample purity was assessed by SDS-PAGE. If not immediately used, the sample was frozen at -20 °C at 1 mM.

**Purification of Aβ42**

Purification of Aβ42 was performed as described previously[4]. A plate of transformed BL21 *E. Coli* with the Aβ42 plasmid was washed and transferred into 200 mL LB and grown at 37 ºC overnight (8-12 hours). The growth was transferred to 1L of LB, grown to $OD_{600}$ of 1.2 and induced with 1 mM IPTG. The cells were resuspended in 40 mL of 6 M guanidinium, 50 mM Tris-HCl pH 8, 300 mM NaCl, and 20 mM imidazole and emulsified with the Avestin Emulsiflex. This lysate was subsequently centrifuged at 23,500 g for 25 min at 4 °C and loaded onto a 5 mL nickel column (HisTrap™, HP, GE Healthcare) equilibrated with 8 M urea, 50 mM NaCl, 50 mM Tris-HCl pH 8, 300 mM NaCl, and 20 mM imidazole. The protein was eluted with increasing amounts of imidazole. The protein containing fraction was loaded on an HPLC at 80 °C and lyophilized. This solid product was then dissolved in 10 mM Tris-HCl pH 6.4, 0.5 mM EDTA, and 1 mM DTT. TEV protease was added in a ratio of Aβ42-fusion:TEV = 6.5:1. This was left until turbid, indicating cleavage had occurred. The solution was centrifuged at 3,500 g for an additional 10 minutes and the pellet was resuspended in 4 mL of 6 M GuHCl, 100 mM NaCl pH 2. This sample was then filtered and reloaded onto the HPLC at 80 °C and lyophilized.

**Growth and purification of SOD**

Codon-optimized MnSOD in the pET21d vector was obtained from GenScript and included an N-terminal 6×His tag, a GST tag, and a TEV protease cleavage site. The plasmid was transformed into *E. coli* BL21 cells and plated. The entire plate was harvested, resuspended in 100 mL of LB medium, and grown at 37 °C to an $OD_{600}$ of 0.8. At this point, the culture was transferred into 1 L of LB medium and grown to a final $OD_{600}$ of 1.2. Protein expression was induced with 1.0 mM IPTG and 8 mM $MnCl_2$, and cultures were incubated overnight (8–12 h) at 20 °C. Cells were harvested by centrifugation at 3,500 g and resuspended in 50 mL lysis buffer containing 100 mM Tris-HCl at pH 8.0, 500 mM NaCl, 10 mM imidazole, 10 mM EDTA, 10 mM BME, 0.2% SDS, and 30% glycerol. One protease inhibitor tablet and 0.4 g streptomycin sulfate were added, and the suspension was incubated for 15 min at 4 °C. Cells were lysed by with an Avestin emulsifier and clarified by centrifugation at 23,500 g. $MgCl_2$ was added to the soluble fraction to a final concentration of 10 mM, and the sample was loaded onto a 5 mL GSTrap column (Cytivia). The column was washed with buffer containing 50 mM Tris-HCl at pH 8.0, 0.5 mM EDTA, and 1 mM DTT until the absorbance returned to baseline. The GSTrap column was then removed from the FPLC system, and TEV protease was manually injected into the column. On-column cleavage was carried out overnight (8–12 h) at room temperature. Following cleavage, the GSTrap column was connected in series to a 5 mL nickel column, allowing elution of MnSOD while retaining TEV protease, free His tags, and uncleaved fusion protein on the columns. The eluted protein was then concentrated to 100 µM and buffer swapped into 300 mM NaCl, pH 7.4, 20 mM KCl, 20 mM magnesium acetate, and 10 mM $MgCl_2$, and frozen at -20 ºC.

**Blue Native Gel Electrophoresis**

Additionally, we aimed to confirm that our purified Hsp10: a cap for Hsp60. Hsp10 was incubated at concentrations of 1 µM in presence of 20 mM KCl, 20 mM magnesium acetate, 50 mM Tris–HCl pH 7.4, 300 mM NaCl, 10 mM $MgCl_2$, 10 mM ATP, and 20 µM Hsp60 for ten minutes at 30 °C. 10 mM mizoribine[5] was added before loading the NativePage gel to stabilize its ATP-dependent state. The NativePage gel was ran at 150 Volts, 110 mAmps, and 120 Watts for 2 hours at room temperature.

**MDH1 refolding assay**

The MDH1 refolding assay was adapted from Gomez-Lorente et al[6]. Briefly, 20 µM MDH1 was denatured by incubation in 10 mM HCl for 30 minutes at room temperature and subsequently diluted to a final concentration of 0.45 µM in 50 mM Tris-HCl (pH 7.4), 20 mM KCl, 20 mM $MgCl_2$, and 5 mM BME, in the presence or absence of 20 µM Hsp10. Samples were further diluted 30-fold with 150 mM potassium phosphate at pH 7.4, 5 mM BME, 1 mM oxaloacetate, and 1 mM NADH and dispensed into Costar 96-well clear flat-bottom microplates in triplicate. NADH oxidation was monitored by measuring absorbance at 340 nm over time at 30 °C using an Agilent BioTek Neo2 multimode reader.

**Calorimetric superoxide dismutase activity assay**

In this assay, xanthine oxidase generates superoxide radicals that react with a 'colorless substrate' to produce a yellow-colored product. The presence of functional MnSOD reduces the accumulation of these radicals, resulting in decreased absorbance at 450 nm. 100 mM MnSOD was denatured with 40 mM HCl and subsequently diluted to a final concentration of 10 mM in a solution containing 50 mM Hsp10. One reaction contained 10 mM functional MnSOD without Hsp10, and one reaction contained 10 mM denatured MnSOD without Hsp10 to show that the MnSOD was adequately denatured. The last reaction contained 10 mM of MnSOD and 50 mM Bovine Serum Albumin (Sigma Aldrich) to act as a negative control to show that the MnSOD was not spontaneously folding alone. Reactions were incubated at room temperature for 5 minutes and then centrifuged at 11,000 g for 5 additional minutes to remove insoluble material. Following centrifugation, 25 µL of xanthine oxidase and 50 µL of substrate were added to each reaction. Each reaction was recorded in triplicates. Absorbance at 450 nm was measured in triplicate over a 30-minute period using an Agilent BioTek Neo2 multimode reader.

**Circular Dichroism**

Circular dichroism (CD) spectroscopy, which reports on secondary structure content in solution, was measured to monitor refolding over time. Since Hsp10 does not contains any α-helical structure, any α-helical signal detected in the spectra can be attributed solely to MnSOD. MnSOD was first denatured at 7 mg/mL in 5 M guanidinium chloride for 10 minutes, followed by incubation with 5 mM EDTA for an additional 10 minutes to chelate the bound $Mn^{2+}$ ion. The sample was then diluted to 0.5 mg/mL in a 20 mM KCl, 20 mM magnesium acetate, 50 mM Tris–HCl pH 7.4, 300 mM NaCl, 10 mM $MgCl_2$, and 5 mM $MnSO_4$. To evaluate refolding, samples were prepared either with or without 0.5 mg/mL Hsp10. CD spectra were recorded after blanking the instrument with either buffer alone or buffer containing Hsp10 to ensure accurate comparison.

**Far-UV circular dichroism secondary structure calculation**

The following formulas were used to determine alpha-helical percentages[7].

$$MRE = \frac{\theta_{obs(mdeg)} \cdot MW}{10 \cdot C \cdot l}$$

$$\%\alpha = \frac{MRE}{-40{,}000} \cdot 100$$

**Mass Photometry**

In order to record concentrations higher than few nM, cover slides were treated according to a protocol by Kratochvil et. al[8]. Cover slides were submerged in acetone and sonicated in a water bath sonicator for 5 minutes. The cover slides were then removed and blown dry with inert gas and submerged in Milli-Q water and sonicated again in a water bath sonicator. The cover slides were removed again and blown dry. The acetone and water bath sonication were repeated for a 2nd time. Next, a 2% v/v APTES solution in acetone was made and heated to 40-50 °C. The cleaned cover slides were submerged and slowly mixed for 30 minutes. The solution was removed from the heat and sonicated for 1 minute, then returned to the heat to mix slowly for 30 additional minutes. The cover slides were removed to a fresh beaker submerged in acetone and sonicated for 5 minutes and left for 1 minute before blow drying. Once dry, a 1 x 1 cm reusable gasket (CultureWell, Grace Bio-Labs) is placed on the center of each cover slide. This well was then filled with 100 µL colloidal silica nanospheres (AlphaNanotech) beneath the water level and incubated for 10 minutes to allow for nanoparticle landing. This solution was gently washed from the gasket with milli-Q water to remove all non-bound nanoparticles. Never was the glass left dry. Next, the water was replaced with 600 mM K2PO4 in 10 mL sodium carbonate/bicarbonate buffer. A 20% w/v PEG solution was made by dissolving 10 mg mPEG-MW5000-SVA in 50 µL of the previous buffer. 50 µL of the PEG solution was then added to each gasket and the coverslips were incubated in the dark for 1 hour to allow for PEGylation of the nanoparticles. After PEGylation, the cover slides were submerged in Milli-Q water and sonicated for 5 minutes with the gasket still attached and subsequently removed and blown dry. After drying, the cover slides were stored at -20 °C in a 50 mL centrifuge tubes with silica packets to prevent moisture. Prior to measurements, a 4-well gasket was placed inside the 1 x 1 cm gasket. All measurements were done on a Refeyn Mass Photometer. Gaussian distributions were determined using the machine-given Refeyn software.

**NMR spectroscopy**

$^{15}N$,$^{2}H$-labeled Hsp10 was used for all Hsp10 titrations recording 2D $^{1}H$–$^{15}N$ HSQC-TROSY spectra in absence or presence of 150 µM MnSOD, 1 mM MDH1, or 500 µM Aβ42. For visualization of the shifts on Aβ42, 60 µM $^{15}N$–$^{1}H$-labeled Aβ42 was titrated with 60–400 µM unlabeled Hsp10. All NMR measurements were carried out in 40 mM sodium phosphate (pH 7), 5% (v/v) $D_2O$, and 0.03% (w/v) sodium azide to prevent microbial growth. The samples had a total volume of 300 µL and were loaded into NMR Shigemi tubes. Experiments using Aβ42 were performed at 277 K, while all other experiments were conducted at 298 K. For both single samples and titrations, the pH was confirmed before and after measurement using an NMR tube–compatible pH probe (Mettler Toledo) to ensure accuracy. All experiments were collected in the Molecular Structures Core, RRID: SCR_022888, in the Department of Chemistry and Biochemistry at the University of Arizona using TopSpin version 4.0 and acquired at 600 MHz on a Bruker Avance NEO spectrometer equipped with a triple-resonance TCI cryoprobe.

NMR data was processed using NMR pipe[9] and analyzed using CcpNMR analysis 3.1[10]. Chemical shift perturbations were calculated as described previously[11] using the equation:

$$\sqrt{\frac{0.125\Delta\delta N^2 + \Delta\delta H^2}{2}}$$

Determination of significance for chemical shift perturbations was performed as described previously[12].

**Thioflavin T assay**

1 mg of purified Aβ42 was suspended in 1.5 mL of 10 mM NaOH and sonicated three times for 30 seconds in a cold-water bath, with 30-second intervals on ice between each sonication. The sample was then ultracentrifuged at 293,000 × g for 1 hour. Samples containing 6 µM Aβ42, 30 µM Hsp10, or a combination of both were prepared in Costar 96-well clear, flat-bottom microplates and monitored with excitation at 450 nm. All samples contained 30 µM ThT in 20 mM KCl, 20 mM magnesium acetate, 50 mM Tris–HCl (pH 7.4), 300 mM NaCl, and 10 mM $MgCl_2$. The plate was read using a BioTek Neo2 multimode reader (Agilent), with fluorescence emission at 490 nm recorded every 5 minutes for 24 hours.

**Cryo-electron Microscopy**

For cryo-EM grid preparation, UltraAufoil R 1.2/1.3 300 mesh Au were glow discharged using a PELCO easiGlow glow discharge cleaning system. Grids were used within 30 minutes from glow discharge was used. Hsp10 was concentrated to 8 mg/mL for cryoEM grid blotting. Grids were prepared using a Leica EM GP2 set to 10 °C and 80% humidity. 4 μL of sample was placed on the grid and after a 10 second wait time, the grid was blotted on Whatman paper and immediately plunge frozen in liquid ethane. A total of 1834 movies were captures using a Titan Krios at 300 kV equipped with the K3 summit detector at the pacific northwest cryo center (PNCC). The movies were measured at 105k magnification with an electron exposure for 50 2-/A$^2$. CryoSPARC was usd to process the movies. 4,317,524 particles were initially picked, with 137,564 particles in the final structure. The final structure has C7 symmetry imposed with a final trdolutin of ~3.7 A.

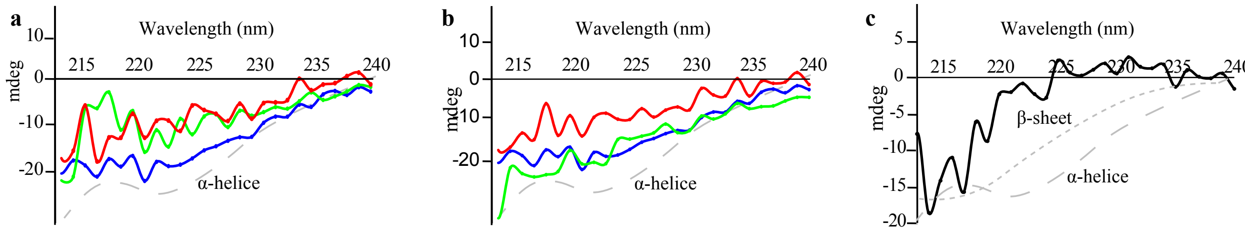


**Supplementary Fig. 1**. A: Far-UV circular dichroism (CD) spectra of native MnSOD (blue), denatued MnSOD (red), and denatured MnSOD with Hsp10 (green) recorded five minutes after dilution. B: Far-UV CD after 15 minutes of incubation of native MnSOD (blue), denatured MnSOD (red), and denatured MnSOD with Hsp10 (green). C: Far-UV CD of Hsp10 alone. MnSOD was denatured at 7 mg/mL with 5 M guanidinium chloride for 10 minutes, followed by incubation with 5 mM EDTA for an additional 10 minutes to chelate the bound $Mn^{2+}$ ion. The sample was then diluted to 0.5 mg/mL in a 20 mM KCl, 20 mM magnesium acetate, 50 mM Tris–HCl pH 7.4, 300 mM NaCl, 10 mM $MgCl_2$, and 5 mM $MnSO_4$. Spectra were either blanked with buffer or buffer with Hsp10, depending on their contents.

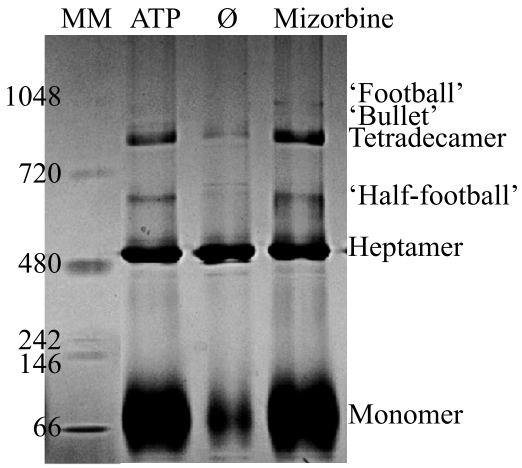


**Supplementary Fig. 2 | Blue Native PAGE demonstrating Hsp10 functioning as a cap for Hsp60.** Hsp60 (20 μM) was incubated with Hsp10 (1 μM) and 10 mM ATP for 10 minutes at 30 °C in buffer containing 20 mM KCl, 20 mM magnesium acetate, 50 mM Tris–HCl (pH 7.4), 300 mM NaCl, and 10 mM $MgCl_2$. Prior to loading onto the gel, the sample was further incubated for 10 minutes with 10 mM mizoribine to stabilize the bound state.

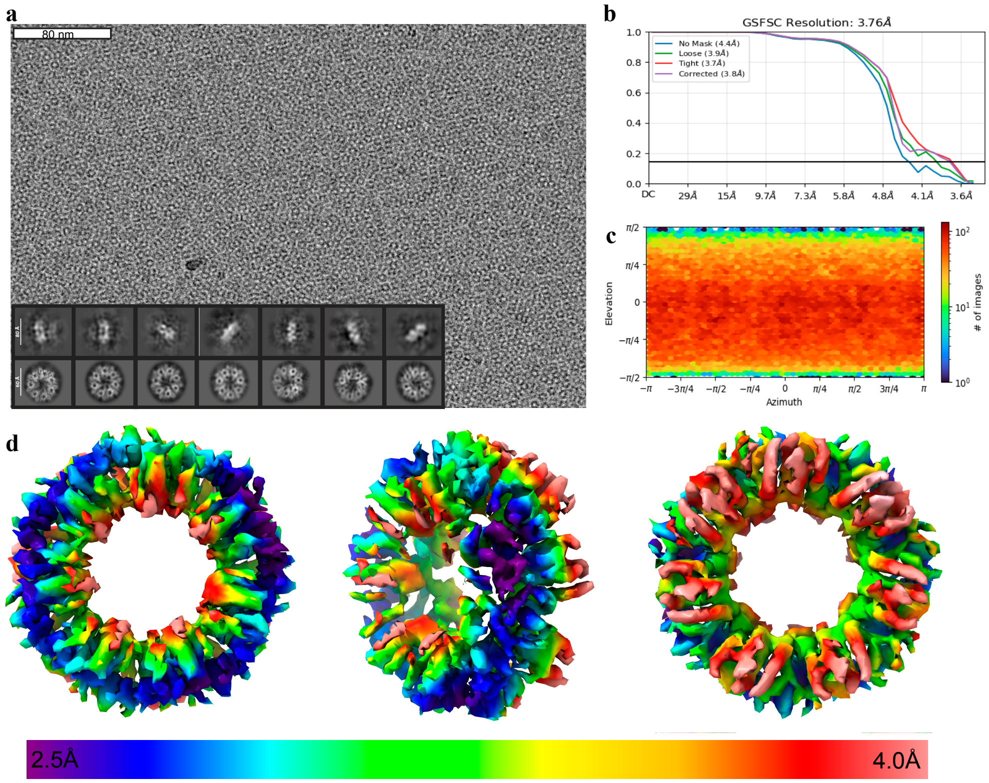


**Supplementary Fig. 3 | Cryo-EM data of double-ring Hsp10. a,** Representative electron micrograph of vitrified Hsp10 at 8 mg/mL, with example 2D class averages from "side" and "top" views. **b,** Fourier shell correlation (FSC) curve for the final 3.7 Å reconstruction generated in CryoSPARC. **c,** Viewing direction distribution of particles contributing to the final 3.7 Å map. **d,** Local resolution estimation calculated in Phenix, colored from 2.5 Å to 4 Å.

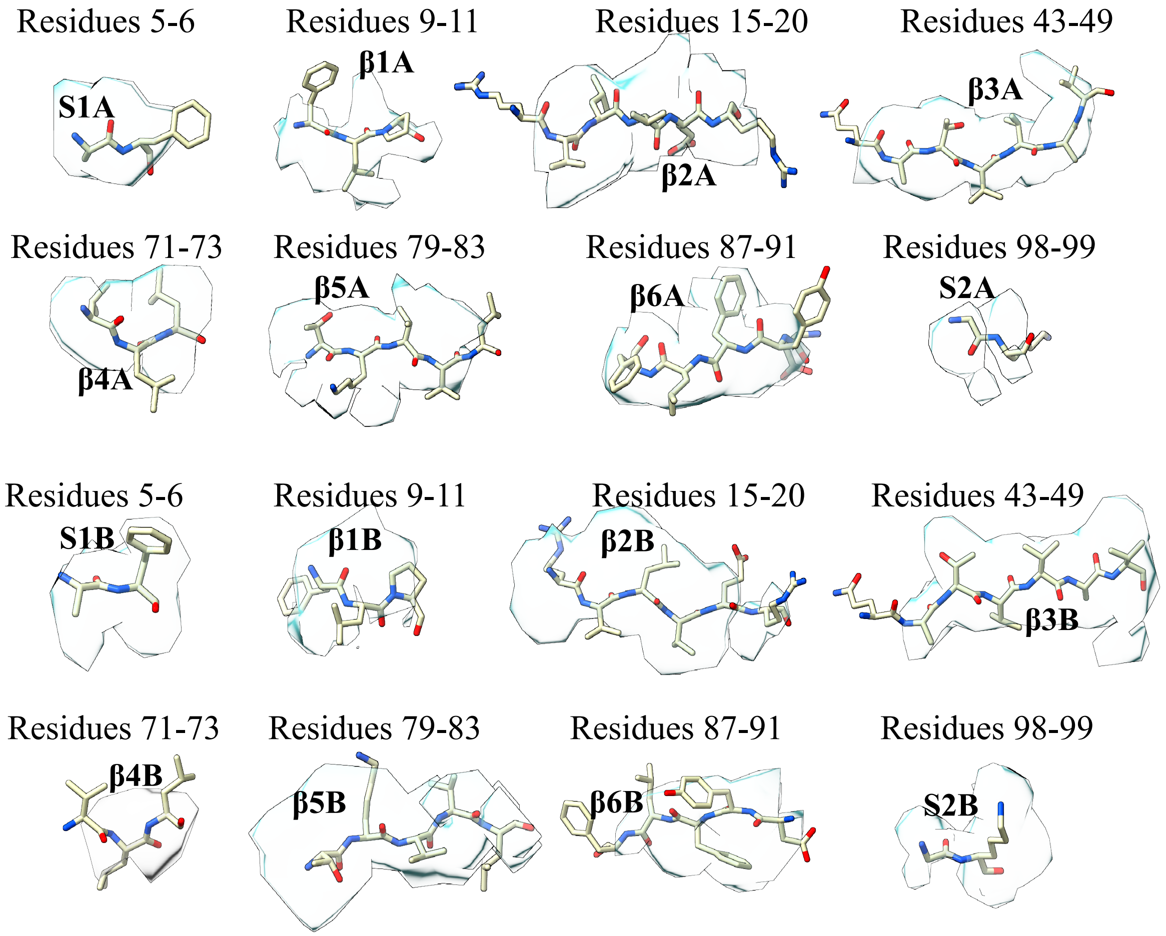


**Supplementary Fig. 4 | Isolated β-strands from the double-ring structure.** The electron density map is shown in transparent mode, with the corresponding β-strands indicated and labeled by their β-strand number (1–6) or segment number (1–2), along with their respective ring (A, B).

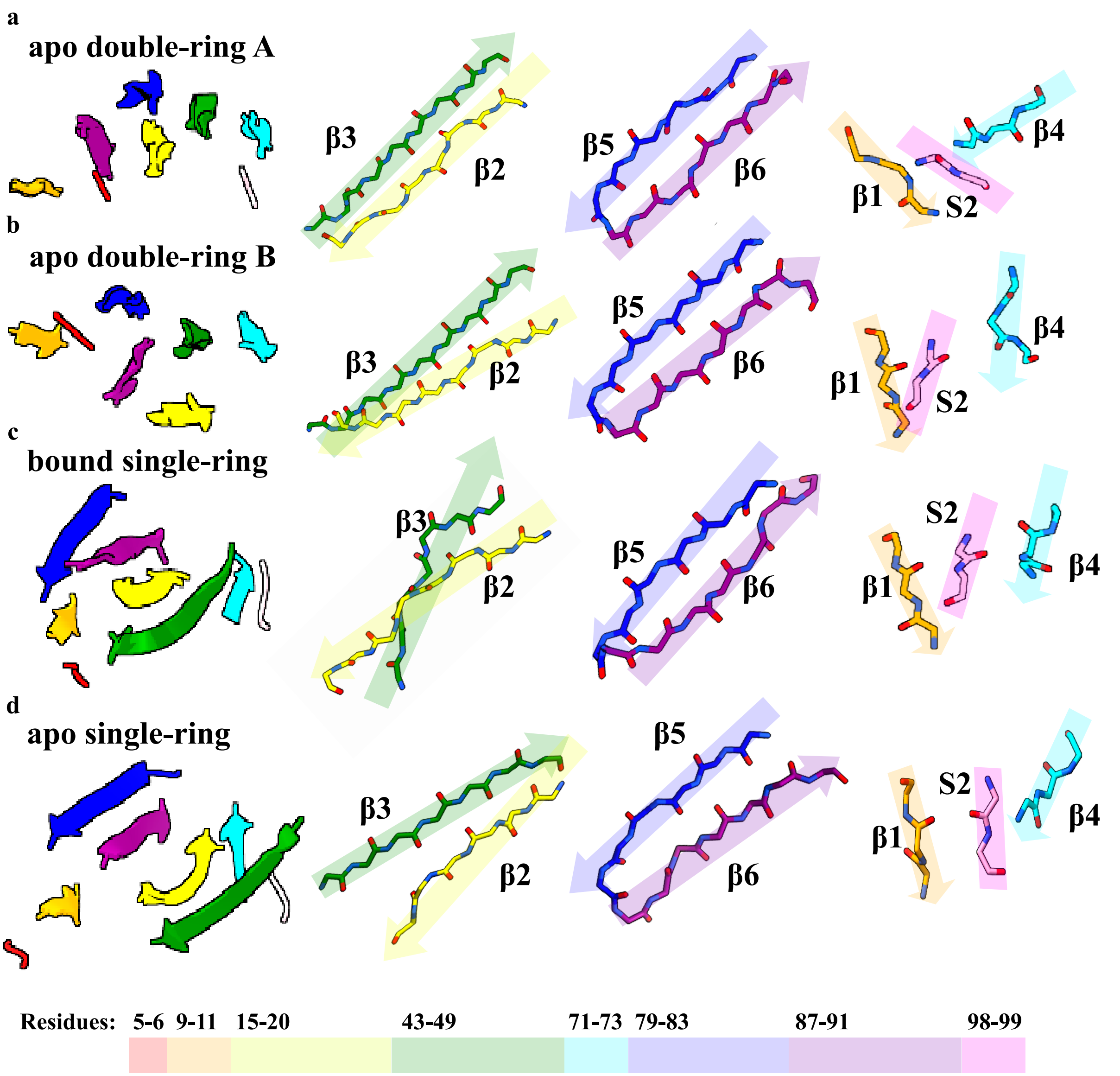


**Supplementary Fig. 5 | Structural comparison of β-sheets in the double-ring and apo single-ring Hsp10, and the published bound single-ring Hsp10 in complex with Hsp60.** Isolated β-sheets comprising β2 and β3 (green, yellow), β5 and β6 (blue, purple), and β1, S2, and β3 (orange, pink, cyan) are shown for the apo double-ring structure (a, b), the bound single-ring structure (c), and the apo single-ring structure (d).

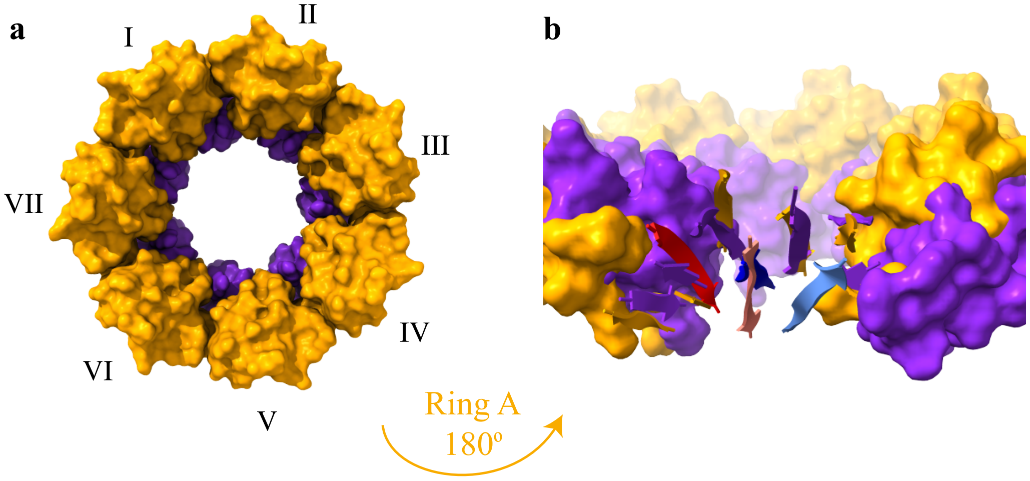


**Supplementary Fig. 6 | Asymmetry of the two rings. a,** The double-ring structure is shown with ring A in orange and ring B in purple; the individual subunits of the heptameric rings are labeled. **b,** Ring A is inverted and overlaid with ring B, revealing structural differences. Notably, two β-strands (β2 and β6) are shifted between the rings: β2 is shown in light blue (ring B) versus blue (ring A), and β6 is shown in light red (ring B) versus red (ring A).

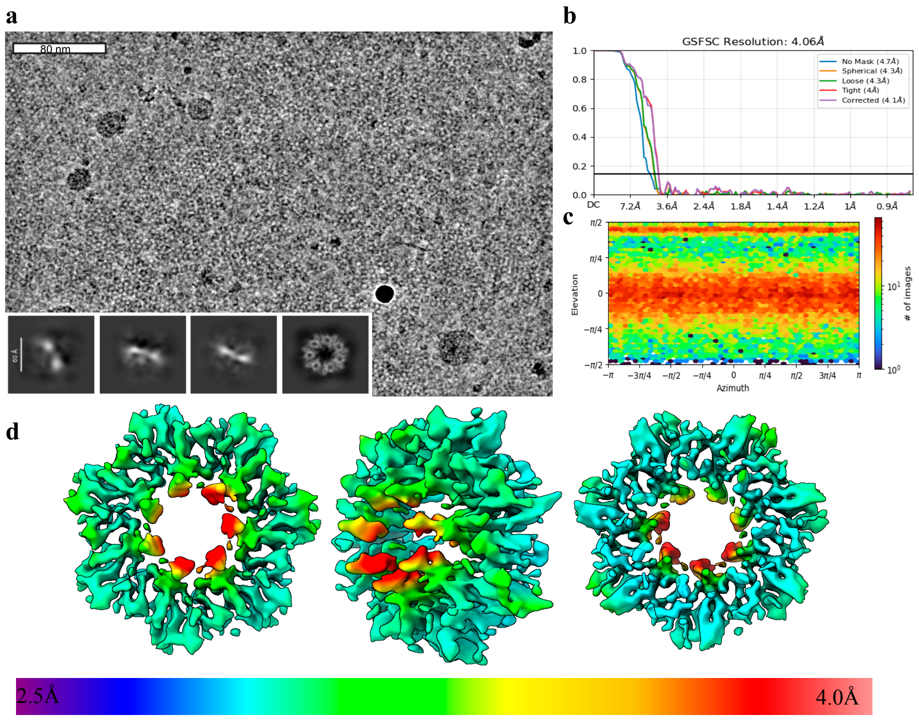


**Supplementary Fig. 7 | Cryo-EM analysis of single-ring Hsp10. a,** Representative electron micrograph of vitrified Hsp10 at 2 mg/mL. **b,** Fourier shell correlation (FSC) curve for the final 4 Å reconstruction generated in CryoSPARC. **c,** Viewing direction distribution of particles contributing to the final 4 Å map. **d,** Local resolution estimation calculated in Phenix, colored from 2.5 Å to 5 Å.

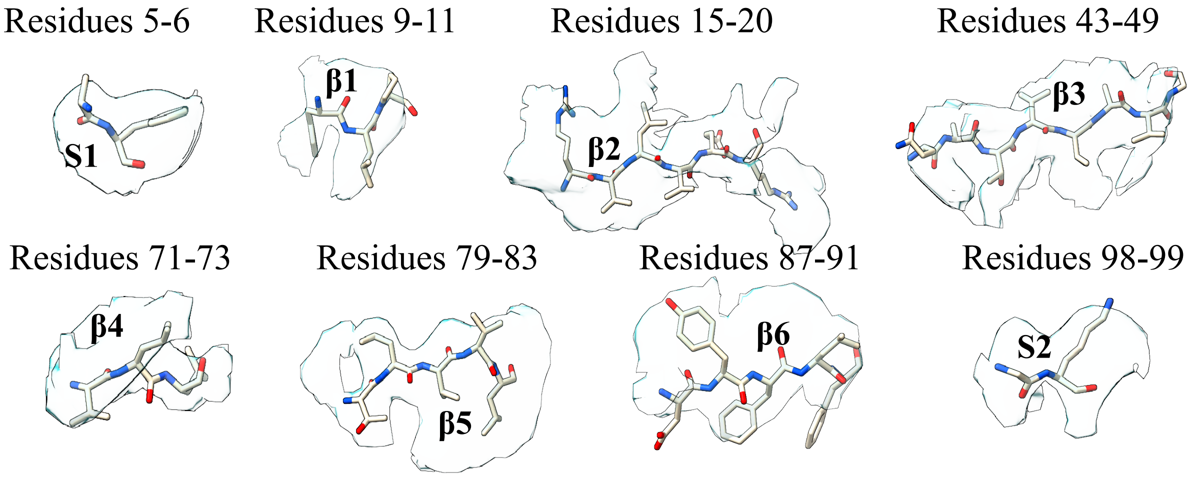


**Supplementary Fig. 8 | Isolated β-strands from the single-ring structure.** The electron density map is shown in transparent mode, with the corresponding β-strands indicated and labeled by their β-strand number (1–6) or segment number (1–2), along with their respective ring (A, B).

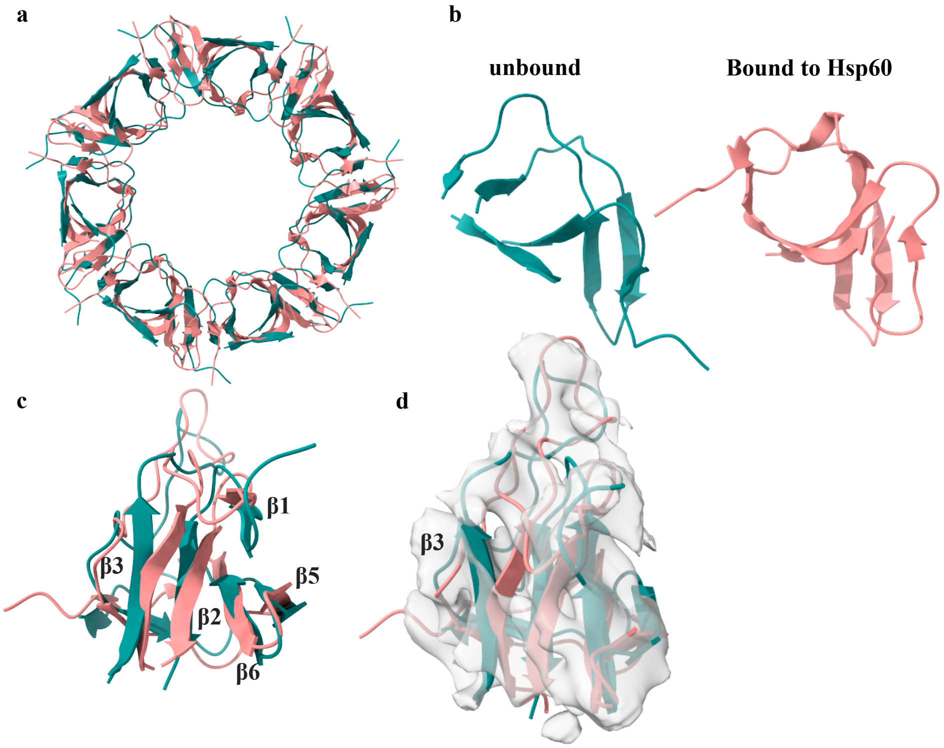


**Supplementary Fig. 9 | Structural comparison of apo and bound single-ring Hsp10**. **a**, Overlay of the apo Hsp10 single-ring structure (teal) with the bound Hsp10 structure (coral) derived from PDB 6MRC. **b,** Top view and **c,** side view of a representative subunit. **d,** Overlay of a single subunit with the corresponding electron density map (gray).

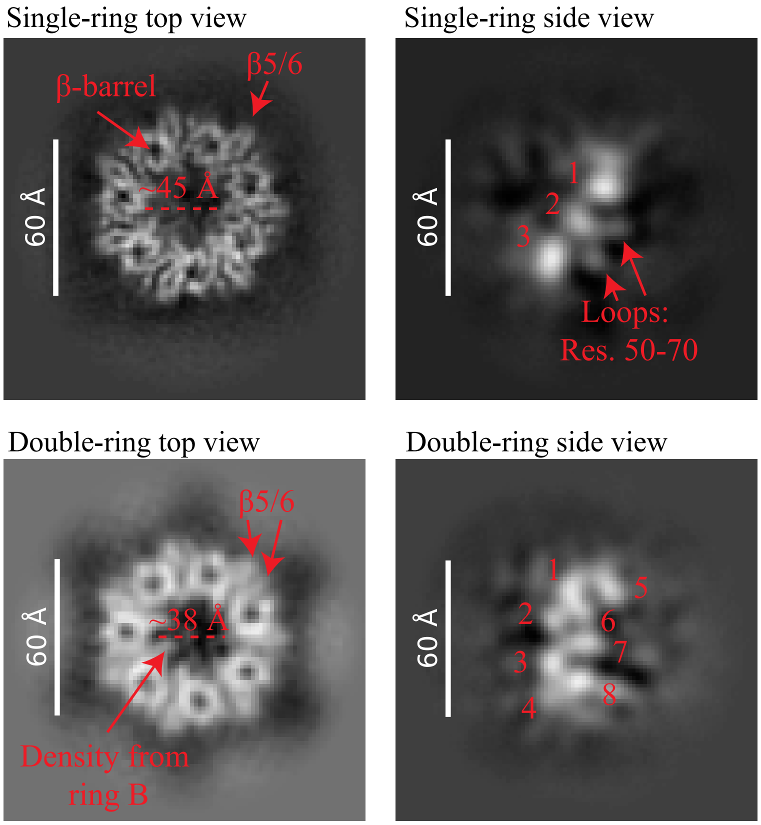


**Supplementary Fig. 10 | Cryo-EM 2D classification of single- and double-ring Hsp10.** Representative 2D class averages of single- and double-ring Hsp10, with domains and key structural features annotated.

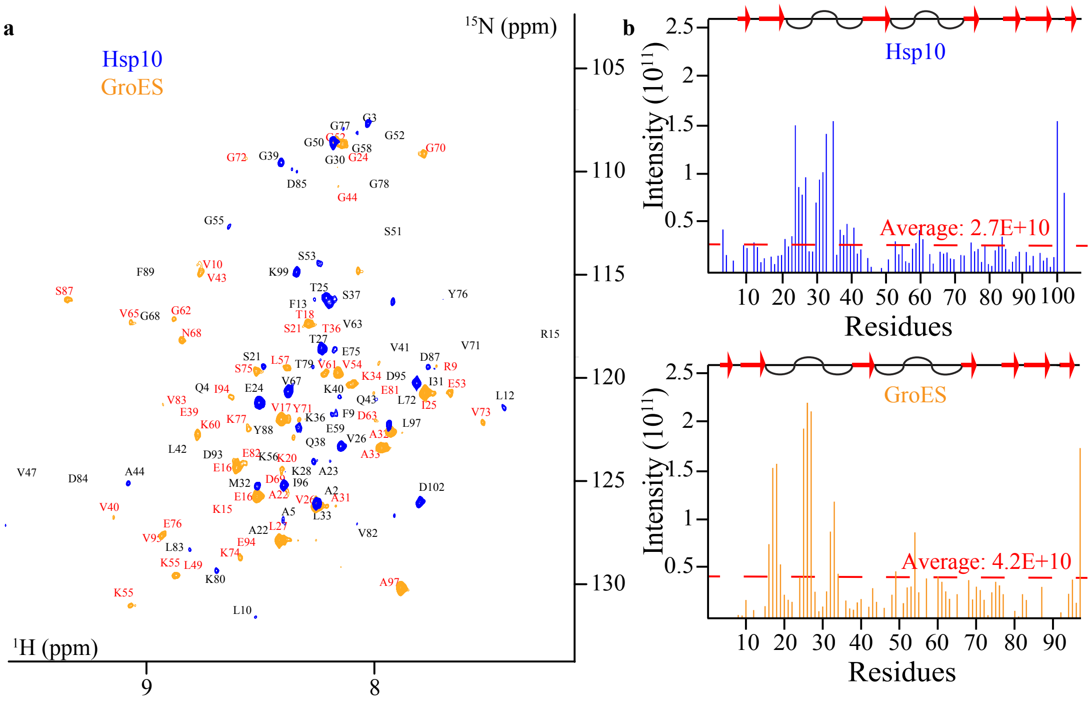


**Supplementary Fig. 11 | HSQC of GroES versus Hsp10. a**, $^{1}H$-$^{15}N$ TROSY spectra of 90 µM [$^{2}H$,$^{15}N$]-labeled GroES (orange) and 90 µM [$^{2}H$,$^{15}N$]-labeled Hsp10 (blue). **b,** Residue-specific intensities for Hsp10 and GroES plotted from panel a. The assignment of Hsp10 and GroES are taken from BMRB accession no. 53226[2], and 7091[13] respectively.

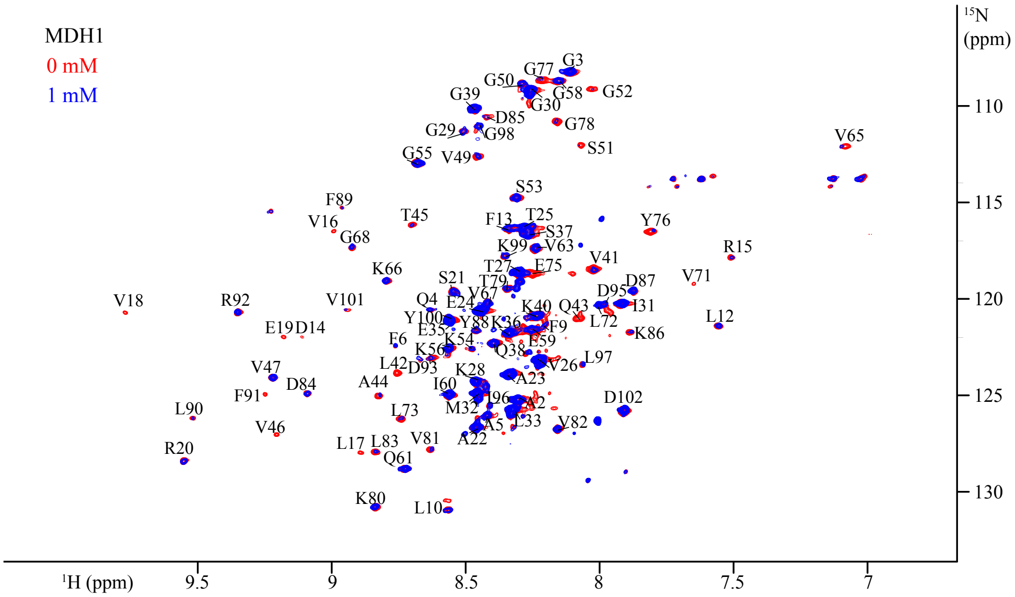


**Supplementary Fig. 12 | 2D $^1H$-$^{15}N$ TROSY spectrum.** 120 µM [$^2H$,$^{15}N$]-labelled Hsp10 in absence (red) and presence (blue) of 1 mM MDH1 was recorded at 600 MHz in 40 mM $NaPO_4$ pH 7.0 buffer at 298 K.

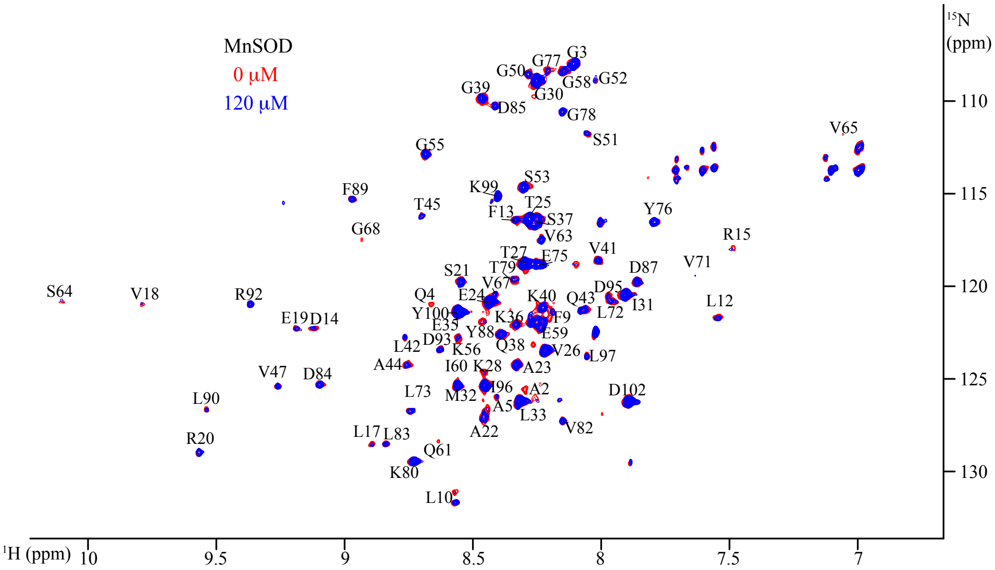


**Supplementary Fig. 13 | 2D $^1H$-$^{15}N$ TROSY spectrum.** 120 µM [$^2H$,$^{15}N$]-labelled Hsp10 in absence (red) and presence (blue) of 120 mM MnSOD was recorded at 600 MHz in 40 mM $NaPO_4$ pH 7.0 buffer at 298 K.

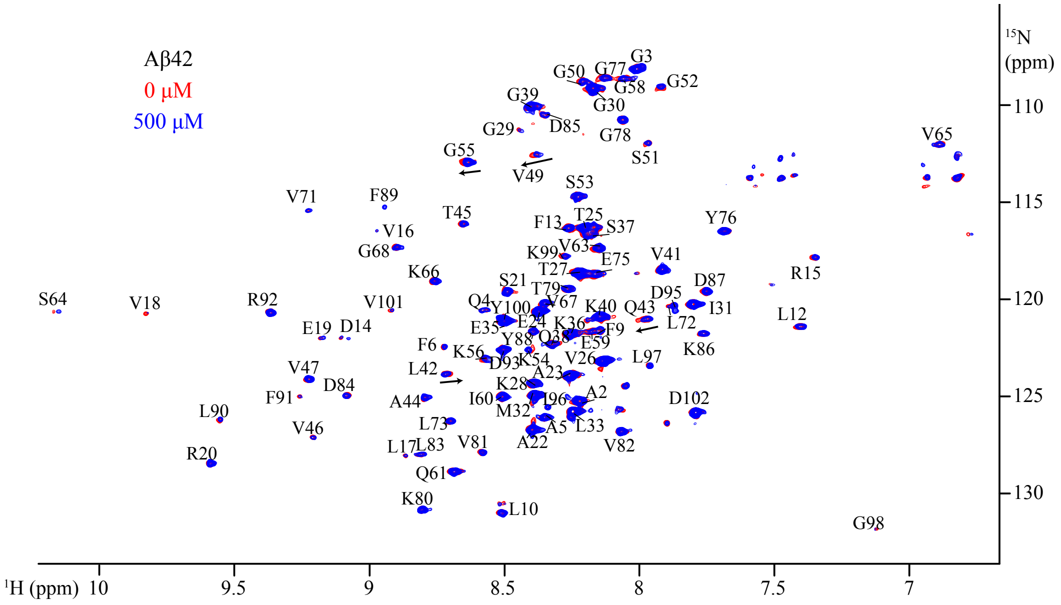


**Supplementary Fig. 14 | 2D $^{1}$H-$^{15}$N TROSY spectrum.** 120 μM [$^{2}$H,$^{15}$N]-labelled Hsp10 in absence (red) and presence (blue) of 500 μM Ab42 was recorded at 600 MHz in 40 mM $NaPO_4$ pH 7.0 buffer at 277 K.

```
Hsp10  11  PLFDRVLVERSAAETVTKGGIMLPEKSQGKVLQATVVAVGSGSKG  55
           PL DRV+V+R   ET + GGI+L   +  K  +  V+AVG+G
GroES  5   PLHDRVIVKRKEVETKSAGGIVLTGSAAAKSTRGEVLAVGNGRIL  49

Hsp10  56  KGGEIQPVSVKVGDKVLLPE-YGGTKVVLDDKDYFLFRDGDILG  98
           + GE++P+ VKVGD V+  + YG     +D+++  +  + DIL
GroES  50  ENGEVKPLDVKVGDIVIFNDGYGVKSEKIDNEEVLIMSESDILA  93
```

**Supplementary Fig. 15 | BLAST sequence alignment of Hsp10 to its bacterial homologue GroES.** Red letters indicated conserved Beta-Sheets, letters between sequences indicate conserved residues, and plus signs (+) indicate conserved chemical environment, but different amino acids

**Supplementary Table 1 |Cryo-EM data collection and processing**

| **Data Collection and Processing** | **EMDB-77112 PDB- 13KK** | **EMDB- 77126 PDB- 13KW** |
|---|---|---|
| Magnification | 105kx | 105kx |
| Voltage (kV) | 300 | 300 |
| Electron exposure (e–/Å$^2$) | 50 | 50 |
| Defocus range (μm) | -0.8 to -2.2 | -0.8 to -2.2 |
| Pixel size (Å) | .8226 | 0.8270 |
| Symmetry imposed | C7 | C7 |
| Initial particle images (no.) | 4,317,524 | 8,872,210 |
| Final particle images (no.) | 137,564 | 46,178 |
| Map resolution (Å) | 3.76 | 4.06 |
| FSC threshold | 0.143 | 0.143 |
| Map resolution range (Å) | 2.5-4.0 | 2.5-5.0 |
| **Refinement** | | |
| Initial model used (PDB code) | None | None |
| Model resolution (Å) | 3.76 | 4.06 |
| FSC threshold | 0.143 | 0.143 |
| Model resolution range (Å) | 2.5-4.0 | 2.5-5.0 |
| Map sharpening *B* factor (Å$^2$) | -100 | -100 |
| **Model composition** | | |
| Non-hydrogen atoms | 3724 | 3913 |
| Protein residues | 406 | 413 |
| Ligand | None | None |
| **R.M.S. deviations** | | |
| Bond lengths (Å) | 1.23 | 0.74 |
| Bond Angles (°) | 1.65 | 1.25 |
| **Validation** | | |

| | | |
|---|---|---|
| MolProbity score | 1.47 | 2.12 |
| Clashscore | 1.95 | 5.77 |
| Poor rotamers (%) | 0 | 0 |
| **Ramachandran plot** | | |
| Favored (%) | 91.18 | 73.91 |
| Allowed (%) | 100 | 100 |
| Disallowed (%) | 0 | 0 |
